# Manipulating prior beliefs causally induces under- and overconfidence

**DOI:** 10.1101/2022.03.01.482511

**Authors:** Hélène Van Marcke, Pierre Le Denmat, Tom Verguts, Kobe Desender

**Author notes:** These authors contributed equally (Hélène Van Marcke, Pierre Le Denmat).

## Abstract

Making a decision is invariably accompanied by a sense of confidence in that decision. Across subjects and tasks, there is widespread variability in the exact level of confidence, even for tasks that do not differ in objective difficulty. Such expressions of under- and overconfidence are of vital importance, as they relate to fundamental life outcomes. Yet, a computational account specifying the mechanisms underlying under- and overconfidence is currently missing. Here, we propose that prior beliefs in the ability to perform a task, based on prior experience with this or a similar task, explain why confidence can differ dramatically across subjects and tasks, despite similar performance. In two perceptual decision-making experiments, we provide evidence for this hypothesis by showing that manipulating prior beliefs about task performance in a training phase causally influences reported levels of confidence in a test phase, while leaving objective performance in the test phase unaffected. This is true both when prior beliefs are induced via manipulated comparative feedback and via manipulating task difficulty during the training phase. We account for these results within an accumulation-to-bound model by explicitly modeling prior beliefs based on earlier exposure to the task. Decision confidence is then quantified as the probability of being correct conditional on these prior beliefs, leading to under- or overconfidence depending on the task context. Our results provide a fundamental mechanistic insight into the computations underlying under- and overconfidence in perceptual decision-making.

## Introduction

Human decision making is accompanied by a sense of confidence regarding the accuracy of those decisions. In experimental work, decision confidence usually correlates with objective accuracy, with participants reporting high confidence for correct choices and low confidence for incorrect choices (1). This tight link between confidence and accuracy is captured in the theoretical proposal that confidence for binary choices reflects the probability of a choice being correct given the available data (2, 3). It follows from this proposal that humans should be rather stable in how they compute and report confidence. However, while decision confidence is on average well explained by such probabilistic models, there exist vast differences between individuals and between tasks concerning the actual reported level of confidence (4). This is clearly visible in simple, low-level perceptual decision-making tasks, where some subjects systematically underestimate and others overestimate their choice accuracy. This phenomenon is often referred to as under- and overconfidence, respectively. Although expressions of under- and overconfidence may not have important consequences in the context of a laboratory task, such biases have far-reaching implications in real life. For example, overconfidence has been related to increased sharing of fake news (5) and diagnostic inaccuracies in physicians (6), while underconfidence has been related to low self-esteem (7). Moreover, impaired insight into the accuracy of one’s decisions (i.e., impaired metacognition) has been linked to holding radical beliefs (8) and a variety of psychiatric symptoms (9).

Despite the clear evidence for individual as well as task differences in confidence, with potentially far-reaching consequences, the origin of these differences is ill understood. Although some researchers have proposed explanations in terms of impression management (10, 11) or exposure to feedback (12, 13), these accounts do not offer a fundamental understanding of the underlying mechanism of biases in confidence. Instead, in the current work we investigated whether biases in confidence can be accounted for within a probabilistic framework. To explain the computational dynamics behind under- and overconfidence, we leveraged an under-appreciated aspect about probabilistic models of confidence: The probability of being correct depends on the task context. Everything else being equal, the probability of a correct choice is higher in an easy than in a difficult task context, simply because correct choices appear more often in easy tasks. Thus, even an agent who simply *believes* to be operating in a difficult task context, will report lower confidence than an agent who believes to be operating in an easy task context (see **Figure 1**). Likewise, an agent who believes that they are very bad at a task, will report lower confidence than an agent who believes they are very competent at performing a task. We introduce the concept of a *subjective drift rate* representing prior beliefs, which controls the mapping between the available data and the probability of being correct. The idea that decision-makers rely on an internal model of the world to inform their computations of decision confidence has been explored before (e.g. (14–17). These modeling efforts have shown that having a “wrong” model of the world could lead to distorted computations of decision confidence. In the current work, we opted for the term “prior beliefs” because within our framework we propose that changes in participants’ beliefs about their own ability to perform a task, i.e., how good they felt they were at a task, influence their computations of confidence. Clearly, having the prior belief that one is good at a task shows clear similarities to having a model of the world that a task will be easy, but the term prior beliefs relates more closely to the idea that confidence depends on participants’ beliefs about their own capacities compared to their beliefs about the world. Apart from theoretical considerations, direct empirical support for the involvement of prior beliefs in the computation of confidence is equally lacking. Our study aimed to provide direct evidence that prior beliefs underlie under- and overconfidence by explicitly manipulating prior beliefs about task performance in perceptual decision-making tasks. In two experiments, we manipulated prior beliefs during the training phase and looked at the influence of this manipulation on confidence ratings during a subsequent test phase. Our results showed that an alteration of prior beliefs, either by means of fake comparative feedback (Experiment 1) or by means of training on tasks with differential difficulty (Experiment 2), selectively affected subsequent (test phase) confidence ratings while leaving performance unaffected. These effects of prior beliefs on decision confidence were accounted for by a probabilistic model of confidence that represented prior beliefs about its ability to perform the task at hand, leading to a different mapping between the accumulated evidence and confidence (see **Figure 1**).

**Figure 1.**
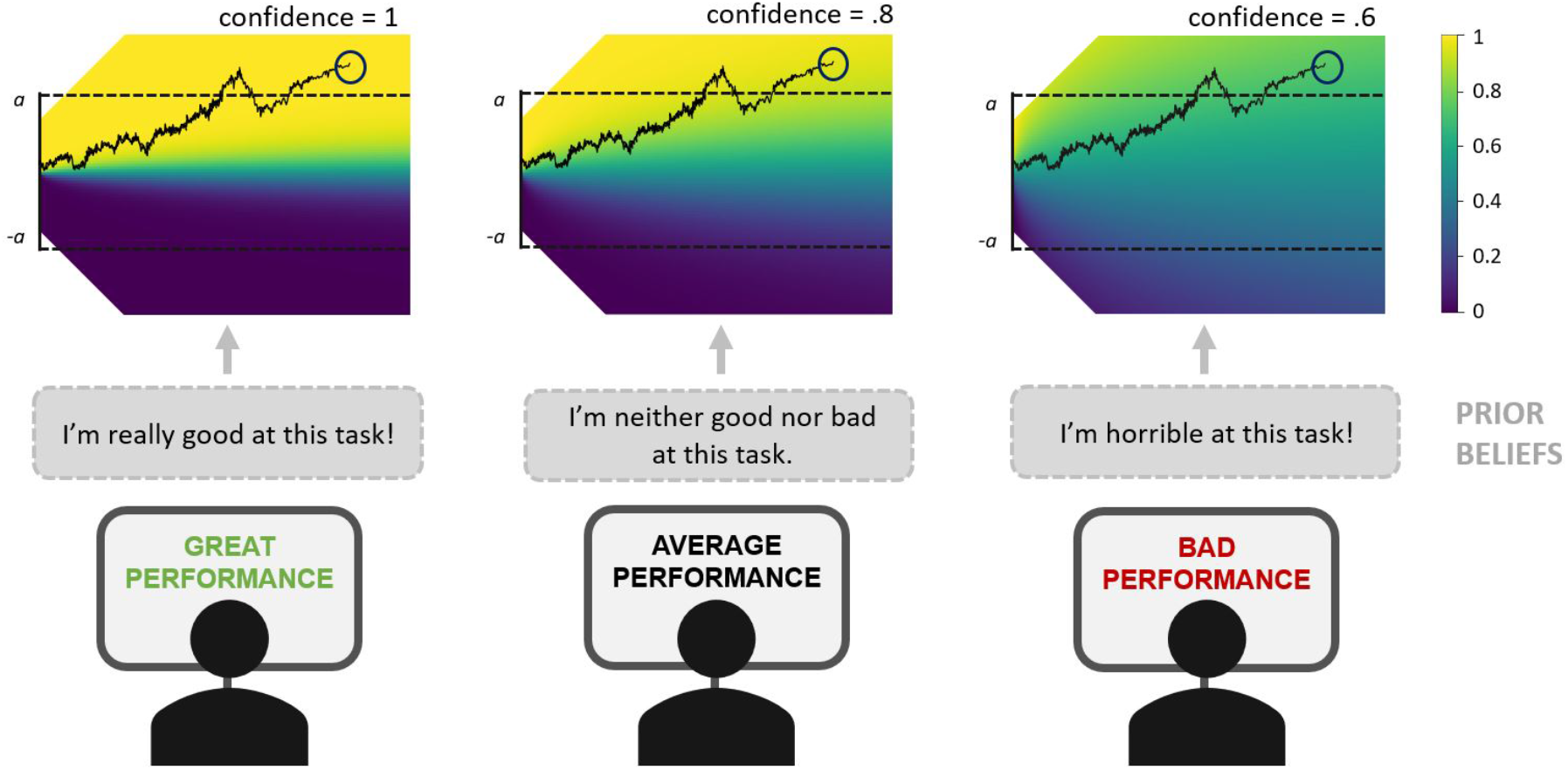
Illustration of how prior beliefs can influence decision confidence. We hypothesized that participants exposed to feedback indicating that they are performing well vs. badly, will hold the prior belief that they are good vs. bad at this task. In our computational framework (visualized at the top), a change in prior belief is implemented by changing the mapping between the amount of evidence (y-axis) and the perceived probability of responding correctly in this task, i.e. decision confidence (colored heatmap). Within this evidence accumulation framework, noisy sensory evidence (y-axis) accumulates over time (x-axis) until one of the two bounds (a or -a) is hit and a choice is made, after which (post-decisional) evidence continues to accumulate and informs decision confidence. In the figure, it can be appreciated that for the exact same trial, the final amount of accumulated evidence (red circle) leads to different levels of confidence depending on the prior belief about task performance.

## Results

To unravel the influence of prior beliefs on decision confidence, we carried out two experiments that aimed to causally influence participants’ prior beliefs about their ability to accurately perform the task. In both experiments, participants performed three similar perceptual decision making tasks. Each task started with a training phase where we manipulated participants’ prior beliefs in their ability to accurately perform the task. This was done by providing them with feedback indicating that their performance was good, average or poor. In the subsequent test phase of each task (without comparative feedback in Experiment 1; without task differences in Experiment 2), we tested the influence of the manipulation on trial-by-trial confidence ratings. To account for the influence of prior beliefs on confidence, we fitted a computational model to the data which holds a belief about its ability to perform the task based on earlier task experience, which can dissociate from its actual performance.

### Experiment 1: Manipulating prior beliefs via comparative feedback causally induces under- and overconfidence

In Experiment 1 (*N*=48), we used comparative feedback to influence prior beliefs about task performance. Participants were told that they would receive feedback every 24 training trials about their performance on the task, relative to a group of participants who had performed the same task at an earlier time. Unbeknownst to participants, feedback was manipulated so that for one task feedback indicated that the participant’s performance was better than most participants’ performance; that it was on average for the second task; and worse than most participants for the third task (see **Figure 2**). Because the feedback was not about performance *per se* but rather about participants’ supposed relative performance, we assumed that the insincerity of the feedback would be noticed less easily. More importantly, we suspected comparative feedback to have a more profound impact on participant’s beliefs about their task performance than direct performance feedback. Afterwards, participants took part in a test phase during which they no longer received feedback but instead rated their perceived level of confidence on each trial. Both during the training and the test phase each task comprised three levels of task difficulty (see Methods). In line with our main hypothesis, confidence ratings during the test phase depended on the feedback that participants received during the training phase, *F*(2,47) = 16.65, *p* < .001, BF_10_ = 5.18e+15 (see **Figure 3C**). Participants reported a higher level of choice confidence after exposure to feedback indicating they performed better (*M* = 4.79), average (*M* = 4.64) or worse (*M* = 4.41) compared to the reference group. This change in average confidence was mostly driven by an increase in “sure correct” ratings and a decrease in “guess correct” ratings after positive versus negative feedback (see **Figure S1, A-C**). Likewise, participants tended to change their mind more often (i.e. reporting guess -, probably - or sure error) after receiving negative compared to positive feedback (see **Figure S1, A-C**). In addition to the effects of feedback, there was the expected effect of trial difficulty on confidence ratings, *F*(2,47) = 159.71, *p* < .001, BF_10_ = 5.21e+51. There was also a small interaction between feedback condition and trial difficulty *F*(4,30744) = 2.60, p = .034, reflecting that the influence of feedback on confidence slightly depended on trial difficulty, but this conclusion was not supported by the BF, which supported the null hypothesis (BF_10_ = .018). As can be seen on **Figure 3C**, the main effect of feedback condition was clearly visible for all levels of stimulus difficulty. Importantly, the induction of prior beliefs selectively affected decision confidence, but left objective performance unaffected. During the test phase, both accuracy and reaction times were affected by trial difficulty (accuracy: χ^2^(2) = 2421.63, *p* < .001, BF_10_ = 9.47e+113; RTs: F(2,30837) = 316.29, p < .001, BF_10_ = 75239472401), but not by feedback condition (accuracy: χ^2^(2) = 0.3, p = .863, BF_10_ = 0.03; RTs: F(2,47) = 2.06, p = .14, BF_10_ = 13.96; see **Figure 3A**). There were also no significant interactions between trial difficulty and feedback condition for objective performance (accuracy: χ^2^(4) = 4.528, p = .34, BF_10_ = .019; RTs: *F*(4, 30837) = 1.024, *p* = 0.3930, BF_10_ = .014). Note that for the effect of feedback condition on reaction times, the BF indicated evidence in favor of the alternative hypothesis. However, this difference seems to originate mostly from the negative condition (whereas the effect of confidence was clearly visible for all three feedback conditions), and was not replicated in Experiment 2.

**Figure 2.**
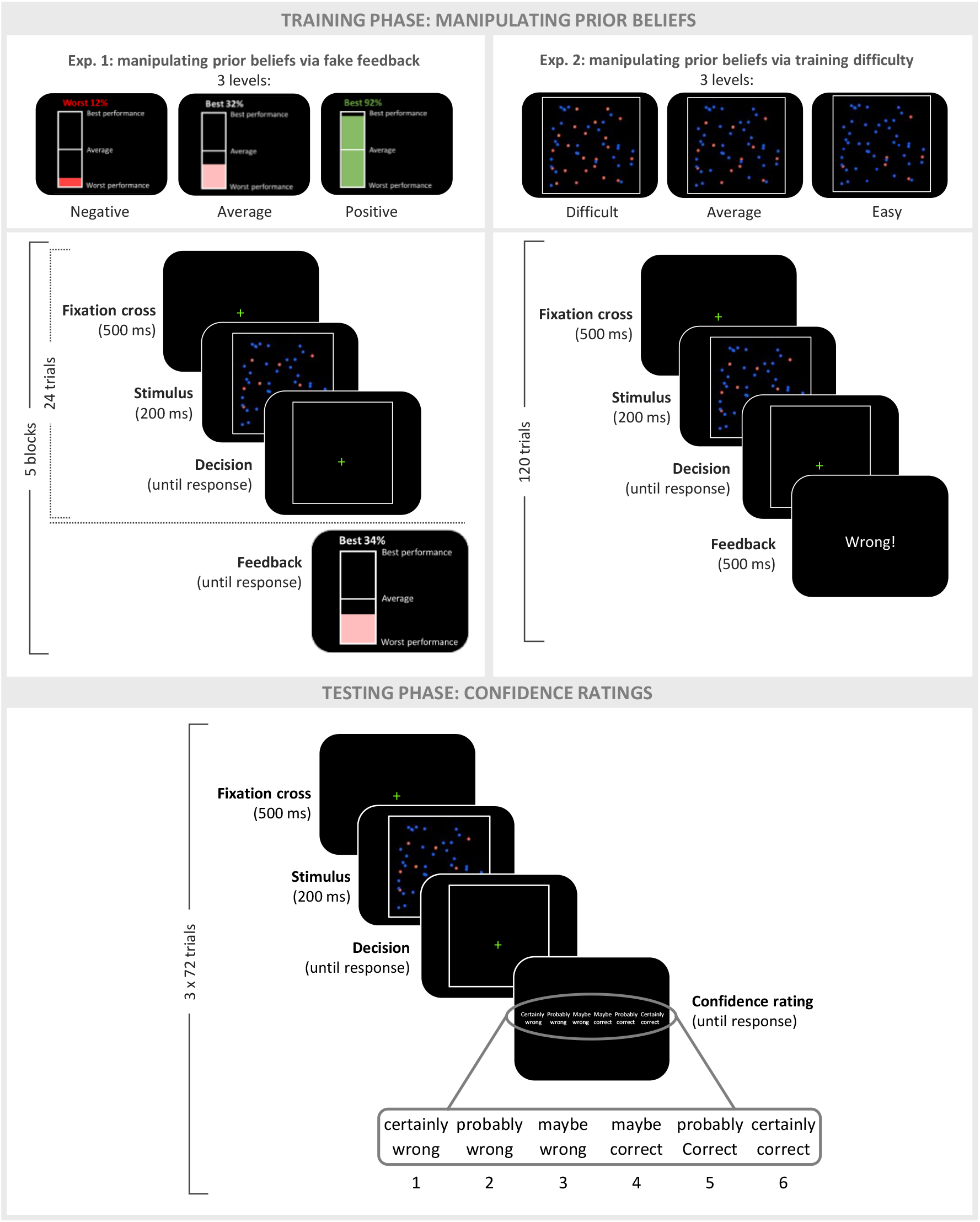
Experimental design. In both experiments, participants performed three different perceptual decision-making tasks (only one shown here). Each task started with a training phase during which a different prior belief was induced. In Experiment 1, participants received comparative feedback after each training block, indicating that their performance was better, similar or worse than the performance of a reference group. In reality, feedback was unrelated to their performance. In Experiment 2, during the training phase participants encountered only easy, average or difficult trials. In this experiment, trial-by-trial feedback reflected their actual performance. For both experiments, these manipulations aimed to install the belief that participants were very good, average or very bad at performing this task, respectively. Each participant was subjected to each of these three manipulations once (i.e., a different manipulation each task). After each training phase, participants completed a test phase during which they no longer received feedback but instead rated their decision confidence after each decision.

**Figure 3.**
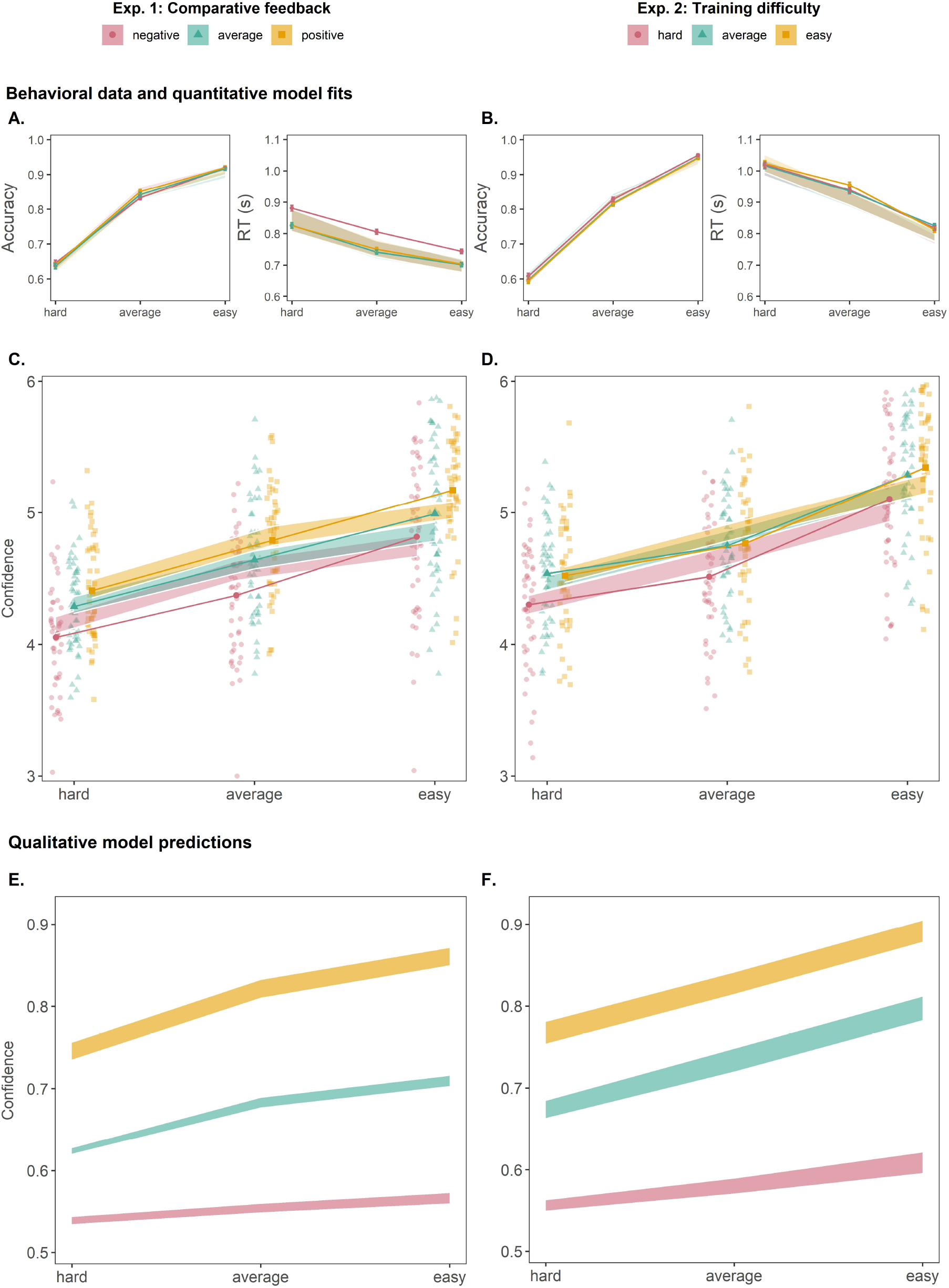
Manipulating prior beliefs causally induces under- and overconfidence. **A**. In Experiment 1 (left column; panels A, C, and E), providing participants with comparative feedback during the training phase indicating that they were performing better, equal or worse than a reference group, left objective performance during the test phase unaffected (A), but induced under- and overconfidence, respectively (C). This effect was captured by our computational model, using both a quantitative (A, C, shaded bars) and qualitative (E) fitting method. **B**. These findings were replicated in Experiment 2 (right column; panels B, D and F), where prior beliefs were manipulated by differential difficulty levels during the training phase. Note: shaded bars reflect the model fits’ standard error of the mean, behavioral data is represented by lines with error bars to reflect standard error of the mean (note that in panels C and D, standard errors were too small to produce visible error bars). Small dots in **C-D** reflect individual participants.

Interestingly, the effect of prior beliefs on confidence was quite persistent throughout the test phase. Each test phase comprised three blocks of 72 trials, separated by a break of one minute. Analyzing the data of each block separately, the effect was remarkably consistent within each of the three blocks (block 1: *F*(2,48) = 21.79, *p* < .001, *M*(positive, average, negative feedback) = 4.79, 4.60, 4.34; block 2: *F*(2,48) = 14.20, *p* < .001, *M*(positive, average, negative feedback) = 4.81, 4.67, 4.44; block 3: *F*(2,48) = 9.51, *p* < .001, *M*(positive, average, negative feedback) = 4.76, 4.65, 4.46). However, there was a subtle decrease in the effect across time: When adding the factor “block” to the main model including the data from all three blocks (see above), there was a significant interaction between block and feedback condition, *F*(4,31412) = 4.98, *p* < .001.

### Experiment 2: Manipulating prior beliefs via differences in task difficulty during training

In Experiment 2 (N=47), we altered prior beliefs about task performance by varying the difficulty of the task during the training phase. Participants were only trained on easy trials on one task, on trials of average difficulty on another task and on difficult trials on a third task (**Figure 2B**). Contrary to Experiment 1, participants received genuine feedback about their choice accuracy (“wrong” or “correct”) on every trial. Critically, because of this difference in difficulty between tasks, we achieved a similar feedback pattern as in Experiment 1: On average, participants saw positive feedback on 95% of the trials (near perfect performance), on 79% of the trials (average performance), or on 60% of the trials (slightly above chance performance) for the easy, average, and hard tasks, respectively. After the training phase, participants took part in a test phase that was identical to Experiment 1 (i.e., three levels of difficulty per task, occurring in equal proportions). Despite the different approach in altering prior beliefs, the results fully replicated those of Experiment 1: Confidence ratings during the test phase depended on the difficulty level of the preceding training phase, *F*(2,46) = 8.19, *p* < .001, BF_10_ = 121461028. Participants reported higher levels of confidence after previous training on an easy task (*M* = 4.88), versus a task of average difficulty level (*M* = 4.86) versus a difficult task (*M* = 4.64; see **Figure 3D**). Similar to Experiment 1, this change was mostly driven by an increase in “sure correct” ratings and a decrease in “guess correct” after training on easy versus difficult trials (for histograms of the actual ratings, see **Figure S1, D-F**). Changes-of-mind (guess -, probably - or sure error) were also more common after training on a difficult compared on an easy task (see **Figure S1, D-F**). As expected, trial difficulty during the test phase also had an effect on confidence ratings, *F*(2,30109) = 2122.11, *p* < .001, BF_10_ = 1.77083e+64, with no interaction between both, *F*(4,30109) = 1.64, *p* = .16, BF_10_ = .02. Again, our manipulation left task performance unaffected. Accuracy and RTs were significantly influenced by testing phase trial difficulty (accuracy: χ^2^(2) = 3090.93, *p* < .001, BF_10_ = 2.013009e+163; RTs: *F*(2, 30109) = 563.52, *p* < .001, BF_10_ = 2.727619e+16), but not by the training phase difficulty conditions (accuracy: χ^2^(2) = .03, *p* = .99, BF_10_ = .09; RTs: *F*(2,46) = 0.01, *p* = 0.99, BF_10_ = .03; see **Figure 3B**). Again, the interaction between both factors was not significant for objective performance (accuracy: χ^2^(4) = 1.6, *p* = .81, BF_10_ = .012; RTs: *F*(4,30109) = 1.52, *p* = 0.190, BF_10_ = .01).

Similar to Experiment 1, the influence of prior beliefs on confidence persisted across time. When adding block to the analysis on confidence reported earlier, there was no significant interaction between training condition and block, *F*(4,30091) = 2.3, *p* = 0.056, and the effect was remarkably consistent across all three blocks (block 1: *F*(2,80) = 15.97, *p* < .001, *M*(easy, medium, difficult training) = 4.88, 4.87, 4.60; block 2: *F*(2,81) = 9.83, *p* < .001, *M*(easy, medium, difficult training) = 4.87, 4.84, 4.65; block 3: *F*(2,82) = 10.69, *p* < .001, *M*(easy, medium, difficult training) = 4.9, 4.87, 4.67).

### Introducing prior beliefs into probabilistic confidence models

In order to address the underlying mechanisms by which prior beliefs influence the reported level of confidence, we turned towards computational models of decision confidence. We focused on accumulation-to-bound models, a family of models that have successfully accounted for choices, reaction time and confidence (3, 18, 19). Accumulation-to-bound models, such as the drift-diffusion model (DDM), describe decision making as the noisy accumulation of evidence until a decision boundary is reached, at which point a response is triggered. The rate of evidence accumulation is controlled by the drift rate (*v*), representing the efficiency of information extraction from the stimulus. To account for decision confidence within such a model, it has been argued that confidence reflects the probability of a choice being correct, conditional on the state of the accumulator (i.e. the amount of evidence accumulated), the decision time and the choice (19–21). In **Figure 1**, this is represented by the heatmaps that visualize how different combinations of evidence (y-axis) and time (x-axis) are associated with different levels of confidence (darker colors are associated with lower confidence). Importantly, when the perceived probability of being correct matches the actual probability of being correct, such a model cannot account for biases in confidence that are independent from objective performance (e.g. such as under- and overconfidence). Intuitively, this occurs because the model’s beliefs about its performance match its actual performance. In a typical evidence accumulation model, task performance is controlled by the drift rate parameter. Importantly, the drift rate also controls the shape of the 2-dimensional heatmap representing probability correct for any given evidence level, time and choice (see **Figure 1**). Thus, higher drift rates will generate heatmaps with a higher probability of being correct than lower drift rates, because high drift rates are associated with higher accuracy and vice versa. To allow for dissociations between actual and perceived performance, we propose that participants have an imperfect approximation of the probability of being correct (which can be manipulated via comparative feedback or differential training difficulty). Thus, we differentiate between *beliefs* about performance and *actual* performance, explicitly incorporating priors beliefs into the computation of decision confidence (for a similar implementation see (16)). In a similar vein, other work has already demonstrated the importance of considering dissociations between participants’ internal model of the world and the external evidence (e.g. (15, 16)). For example, Khalvati et al. (2021) were able to show that common discrepancies between confidence and choice accuracy can be explained by assuming a wrong model of the world. Although Khalvati et al. used a Bayesian framework, the similarity between DDM and Bayesian models has been established (22). Formally, we propose to parameterize the computation of the probability of being correct and thereby provide a solution as to how individuals integrate previous experience with the current task to form prior beliefs about current performance. To achieve this, we propose a dissociation between the drift rate controlling objective task performance, and the *subjective* drift rate controlling the shape of the heatmap (i.e. representing probability correct). This *subjective* drift rate can be thought of as a formalization of prior beliefs (inverting the heatmap into a single parameter), reflecting how good participants *think* they perform at a task rather than how they actually perform (see Methods for full details). Thus, different values for the subjective drift rate will give rise to different, unique probability maps, corresponding to different, unique prior beliefs. By assigning different values to the subjective drift rate while leaving the other parameters of the model unaffected, this proposal can in principle explain how conditions with identical objective task performance (i.e., same drift rates) but different prior beliefs (i.e. different subjective drift rates) can lead to differences in subjective confidence. That is precisely the pattern of behavior observed in both experiments: In Experiment 1, participants were faced with false comparative feedback in the training phase, in the sense that it misinformed them about the positioning of their task performance relative to the performance of others. In Experiment 2, since participants were exposed in the training phase to only one of the three difficulty levels subsequently experienced in the test phase, they received an accurate, yet necessarily biased sample of the heatmap.

### Modeling the effect of prior beliefs on decision confidence

#### Quantitative model fitting

To validate the prediction of our model that differences in subjective confidence, but not task performance, can be captured by a change in subjective drift rate only, we fitted our model to the performance and confidence ratings observed in the testing phase. Given that there was no effect of the comparative feedback or training difficulty manipulations on performance, we estimated DDM parameters (bound, non-decision time and drift rate) based on testing phase RTs and accuracy data separately for each participant, with fixed DDM parameters over conditions. Subjective drift rate was estimated separately for each participant and prior belief condition to the confidence ratings in the testing phase. Model-predicted confidence was discretized into partitions of equal intervals in order to be mapped on the same 6-point scale as the confidence ratings. We fitted an additional bias parameter to account for the specific mapping from continuous probabilities to the categorical ratings that participants made. Importantly, this parameter was fixed over conditions, so that each participant only has one bias parameter. As can be seen on **Figure 3, A-D**, model fits closely tracked empirical accuracy, RTs and confidence ratings. Importantly, simulated confidence ratings from the best-fitting parameters showed the same pattern as the empirical data. Simulated confidence ratings increased both with increasingly positive feedback presented in Experiment 1, *F*(2,48) = 6.92, *p* = .002, BF_10_ = 7.4838542e+7, and with easier training difficulty in Experiment 2, *F*(2,47) = 7.02, *p* = .002, BF_10_ = 1.93840824e+8. Simulated confidence ratings were also influenced by trial difficulty in both Experiment 1, *F*(2,48) = 194.27, *p* < .001, BF_10_ = 8.145035e+36 and Experiment 2, *F*(2,47) = 251.68, p <.001, BF_10_ = 1.978466e+61. Lastly, identical to behavioral data, no interaction was found between prior belief condition and trial difficulty (Experiment 1: *F*(4,30306) = 1.07, *p* = .37, BF_10_ = .01; Experiment 2: *F*(4,29519) = 0.38, *p* = .82, BF_10_ = .01).

#### Qualitative model fitting

In the previous section, we demonstrated that our computational model was able to capture the influence of prior beliefs on confidence by assuming a change in subjective drift rate between the different conditions. We next show that our model can also account for the influence of prior beliefs on confidence even when it is blind to empirical confidence ratings. In this section, our model was exposed to the same training conditions as participants, and was then asked to predict confidence judgments based on the performance in testing phase. For this sake, we estimated the subjective drift rate, per participant and per task, based on the data of the training phase. To do so, we estimated DDM (23) parameters based on the training phase data and generated simulations based on these parameters. We estimated which subjective drift rate parameter provides confidence predictions that are in line with the feedback presented to participants. As expected, when the model was exposed to negative feedback, the estimated subjective drift rate was lower compared to when it was exposed to positive feedback, *F*(2,94) = 450.02, *p* < .001 (Experiment 1, **Figure 4A**). Likewise, when the model was trained on a difficult task, the estimated subjective drift rate was lower compared to when the model was trained on an easy task, *F*(2,92) = 64.97, *p* < .001 (Experiment 2, **Figure 4E**). Second, to demonstrate that our feedback- and training manipulations selectively influenced subjective drift rate but left objective performance unaffected, we next estimated the parameters of our accumulation-to-bound model based on the test phase data as well. The estimated parameters did not vary with the feedback conditions in Experiment 1 (all *p*s > .36, **Figure 4B-D**), nor were they influenced by the differential training difficulty in Experiment 2 (all *p*s > .31, **Figure 4F-H**). Thus, our model was able to generate different levels of prior beliefs about task performance after seeing fake comparative feedback (Experiment 1) or performing tasks of differential training difficulty (Experiment 2).

**Figure 4.**
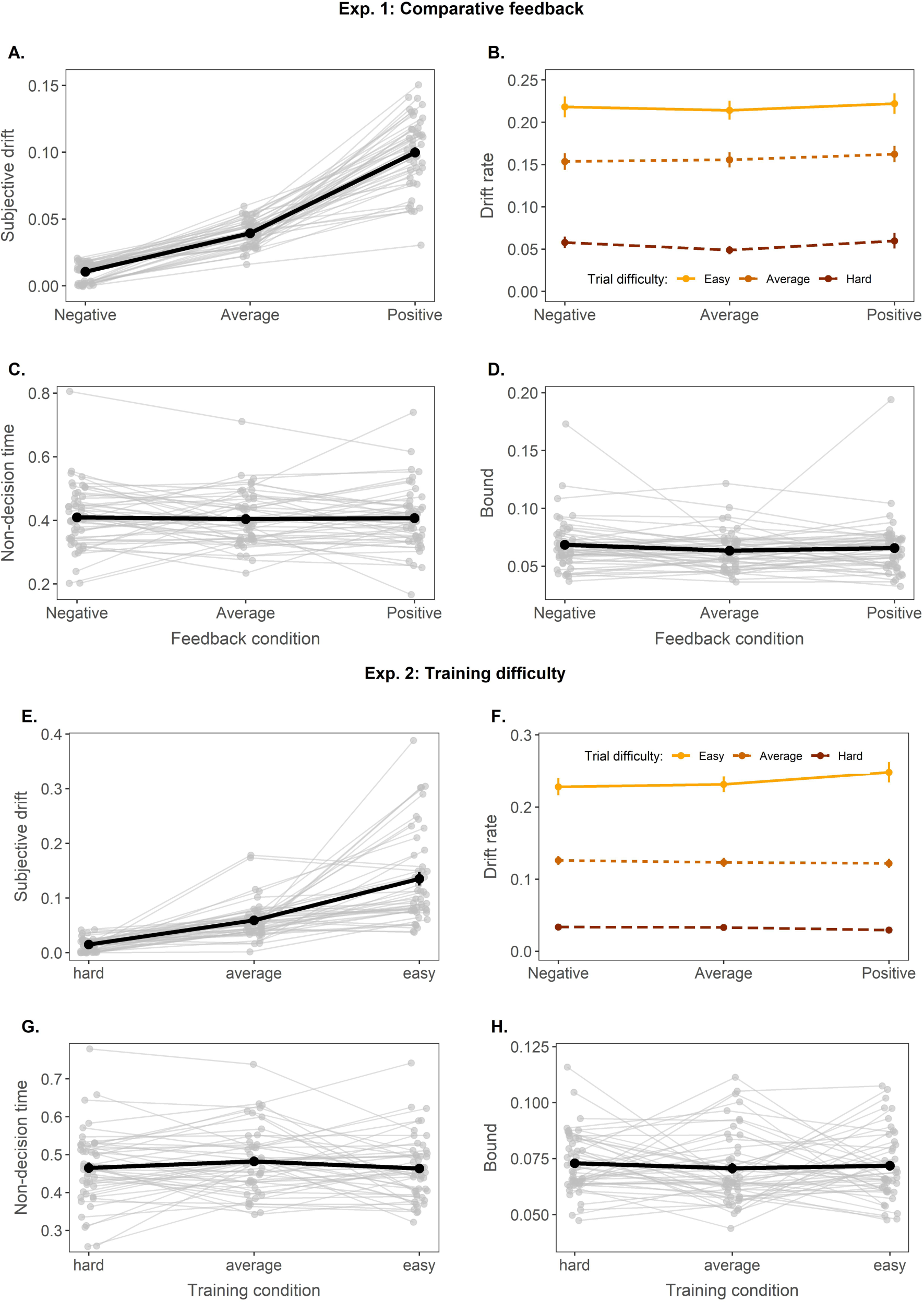
Manipulating prior beliefs selectively influences subjective drift rate. **A.** In Experiment 1, the subjective drift rate (reflecting the prior belief about performance) increased when the model was exposed to increasingly positive feedback. **B-D**. The feedback conditions in the training phase did not influence the other parameters of the evidence accumulation model. **E-H**. Similarly, in Experiment 2, only the subjective drift rate was sensitive to the differential training difficulty. Note: grey, dotted lines reflect individual participants, solid black lines represent the mean, error bars reflect standard error of the mean.

Third, we finally tested whether this difference in prior belief induced during the training phase was sufficient to capture under- and overconfidence in the test phase. To do so, we queried model predictions based on the DDM parameters obtained from the fit to the data of the test phase, but using the subjective drift rate which was estimated from the data of the training phase. It is important to stress that using this approach, instead of fitting our model to empirically observed confidence data, we instead generated model predictions from a model that was merely exposed to the same feedback as the participants. Thus, our model was effectively blind to the empirical confidence judgments. As expected, for both Experiment 1 and Experiment 2 (**Figure 3E**) the model predicted increases of confidence with increasingly positive feedback (*F*(2,47) = 274.10, *p* < .001, Experiment 1), and lower task difficulty during the training phase (*F*(2,46) = 91.00, *p* < .001, Experiment 2). Additionally, the model also predicted the expected increase of confidence with lower testing trial difficulty, (Experiment 1: *F*(2,282) = 233.32, *p* < .001; Experiment 2: *F*(2,276) = 353.67, *p* < .001). Finally, for both experiments there was an interaction between both factors, *p* < .001, reflecting that the model predicted the effect to be slightly smaller with difficult trials. In sum, we successfully accounted for expressions of under- and overconfidence within accumulation-to-bound models by taking into account prior beliefs.

## Discussion

The level of confidence expressed in choices differs greatly between humans and between tasks. These differences in the overall level of confidence are often referred to as under- and overconfidence. Here, we provide direct evidence that, for perceptual decisions, these differences arise from prior beliefs about the ability to perform a task. In two experiments, we have shown that a manipulation of prior beliefs causally influences the reported level of decision confidence. Furthermore, we demonstrate that this finding can be accounted for by extending probabilistic models of confidence with a *subjective* drift rate, explicitly representing prior beliefs about task performance within these models. Our behavioral manipulation of prior beliefs selectively influenced the model’s prior belief, which in turn accounted for under -and overconfidence as seen in the data.

### The mechanism behind under- and overconfidence

The last decade has seen an increase in studies investigating how the sense of confidence can be explained within models of decision making. However, while most prior work has focused on measures that quantify the sensitivity of decision confidence (15, 24, 25), much less attention has been devoted to the computational mechanisms underlying confidence biases. For example, in signal detection theory (26, 27), biases in confidence can easily be modeled by changing the criteria that dissociate high from low confidence (28). However, this is merely descriptive and does not provide us with fundamental insight into the computational mechanisms underlying under -and overconfidence. To answer this question, in the current work we relied on accumulation-to-bound models that explain confidence as the posterior probability of being correct given available data. Specifically, we consider both time and evidence as relevant data to compute this probability (3, 18). Although it has been suggested previously that prior experience might be an important factor to understand deviations in the computation of confidence (16, 17, 21), empirical evidence for this claim has been lacking so far. Here, we provide the first empirical demonstration that inducing under- and overconfidence by means of changes in prior beliefs can be readily accounted for within dynamic probabilistic models (see (29) for an explanation in terms of expected precision). Moreover, we demonstrate this using both a quantitative and qualitative fitting method, i.e., fitting our model directly to the empirically observed confidence ratings to query quantitative predictions as well as merely exposing the model to the same training conditions that participants were presented with and querying qualitative predictions. Both fitting methods successfully captured the patterns we observed in the empirical data, predicting an effect of comparative feedback as well as different training difficulties on subsequent confidence ratings. Our quantitative fitting method provided a close fit to the empirical data, and captured subtle patterns in the data such as the close similarity between the easy and medium condition in Experiment 2. In contrast, our qualitative method predicted stronger effects than those actually observed and additionally predicted an interaction between training condition and testing phase difficulty that was not present in the empirical data. Still, it is not trivial that a model that was effectively blind to the empirical confidence judgements, successfully captured the empirical finding that the reported level of decision confidence depends on the training phase conditions that participants were exposed to. An important consideration here is that during the qualitative fitting approach, our model starts from a blank slate (i.e. without any prior knowledge or preconceptions) and then builds its prior beliefs entirely based on the feedback. Real participants, however, likely come to the experiment with pre-existing prior beliefs about their abilities, and our experimental manipulations ride on top of these preconceptions. As such, a fruitful approach for future work will be to unravel how our experimental manipulations interact with earlier prior beliefs about task performance. As a second note, our model assumed perfect integration of the feedback. This might be different from real humans, who likely display leaky integration of feedback. Nevertheless, despite these simplifications, the qualitative predictions of our dynamic probabilistic model were in line with the empirical data. Although our prior belief induction is necessarily lab-based and thus slightly artificial, it is not hard to imagine how this process might operate in real life. Spontaneous differential exposure to comparative feedback (cf. Experiment 1) or mostly engaging in difficult versus easy tasks (cf. Experiment 2) will result in differences in prior beliefs about task ability and hence confidence ratings.

### The interplay between local and global confidence

The findings reported in the current work are closely linked to the formation of so-called global decision confidence, and the way in which this global confidence can influence the computation of local confidence judgments. In recent years, there is growing interest in the computation of global decision confidence, that is confidence at the task level (30, 31). Global confidence is the general, subjective feeling that subjects have about their ability to perform a task, spanning a broader timescale than trial-specific local confidence. From this description, it becomes clear that global confidence bears close resemblance to the concept of prior beliefs and the task-general subjective drift rate discussed in the current work. Following this logic, our findings thus suggest a direct influence of prior beliefs about task performance (global confidence) on how we actually believe we are doing on individual trials (local confidence). Interestingly, it has been shown that in the absence of trial-by-trial feedback, subjects compute global confidence by integrating local confidence judgments (30). Thus, there also seems to be a direct influence of local confidence on global confidence, revealing an intriguing interplay between both. Therefore, it could be that causally inducing prior beliefs might have a long-lasting, self-sustaining effect on local confidence by means of a self-sustaining loop between local and global confidence. Our current data already suggest that the effect of the prior beliefs manipulation is not limited to the first few test trials only: Even in the third, final test block (72 trials), the effect was still visible, suggesting a long-lasting effect on confidence instead of a mere temporary boost or lapse in self-confidence specific to the training phase only. Future work might address whether the persistent nature of prior beliefs on local decision confidence is indeed mediated by global confidence. As impaired confidence has been linked to a variety of psychiatric symptoms (9), uncovering the mechanisms behind these persistent biases could provide important new insights for clinical practice. For example, the self-sustaining nature of our prior beliefs manipulation could potentially provide a new tool to aid in restoring impaired confidence estimation in individuals with psychiatric disorders.

### Dissociations between accuracy and confidence

Although confidence generally tracks objective accuracy (32), an increasing number of recent studies have reported dissociations between confidence and accuracy (28, 33). For example, it has been shown that whereas choices are equally informed by choice-relevant and choice-irrelevant information, decision confidence has been found to mostly reflect variation in choice-relevant information (34–37). In a similar vein, it has been reported that variance has a more profound effect on confidence than it has on decisions (38–41). Importantly, these observations are often treated as evidence for the existence of a metacognitive module existing separately from the decision making circuitry (15, 42). In the current work, we reported a clear dissociation between accuracy and confidence, with only the latter being influenced by our manipulations of prior beliefs. Importantly, different from earlier work, our interpretation of these findings does not require a separate metacognitive processing stream during the computation of confidence. Instead, we were able to explain decision confidence within the decision circuitry by simply changing the prior beliefs within this framework (16). One could argue, still, that the process of forming (and updating) prior beliefs about task performance as described in our model is the work of a metacognitive module. However, in contrast with those modules described in earlier works (15, 42), our module is rooted within the decision-making process. In other words, our model does not assume processing of metacognitive evidence independent and parallel to the processing of sensory evidence. All in all, our findings add to the ongoing debate about the need for a separate metacognitive module to explain dissociations between accuracy and confidence, demonstrating that both can in fact operate on the same stream of data.

### Counterfactual confidence

One interesting discussion point is the extent to which participants were aware that they were reporting different levels of confidence for the same levels of evidence. Especially in Experiment 1, 18 out of the 49 participants indicated in the post-experiment debriefing that they were aware of the influence the comparative feedback had on their confidence ratings. These participants explicitly stated that positive feedback made them feel more confident about their answers, while negative feedback had the opposite effect. Notably, an additional analysis showed no difference between these and the other participants in terms of our manipulation’s effect on confidence. This raises an intriguing question about whether participants immediately computed the level of confidence that they eventually reported (i.e., modulated by prior beliefs), or rather whether they initially computed the “unbiased” probability of being correct and strategically lowered or increased this rating depending on the feedback seen earlier. The latter option would imply that participants have an unbiased representation of confidence at their disposal, which could be used for alternative purposes. Just like the representation of confidence based on external feedback that we described with our model, this unbiased representation could be formed in a similar way from an unbiased internal feedback signal. Although such a dissociation between a biased and unbiased confidence estimate might sound unlikely, previous work has shown evidence that participants compute so-called “counterfactual” confidence which they use to update their response bias (43). Moreover, in social contexts it is known that people can sometimes feel (not) very confident but for social reasons decide to report a (higher) lower level of confidence (44). Such modulations of reported confidence could be accounted for by our model by considering the decision-maker holds a more explicitly aware bias between the actual computation and the reporting of confidence.

### Suggestions for future research

In the current work, we show that under- and overconfidence can be explained as resulting from prior beliefs about the ability to perform a task. This hypothesis was tested in two separate experiments making use of perceptual decision-making paradigms. To further confirm and solidify the generality of our claim, future research should focus on the role of prior beliefs in other domains than perceptual decision-making (e.g., memory, learning, etc.) and finally, test this hypothesis in real-life decision making (e.g., economic decision-making). Our manipulation of prior beliefs through comparative feedback (Experiment 1) could be particularly fruitful to study social aspects of confidence in group decision-making. While in Experiment 1 we compared participants’ performance to that of so-called other participants, participants completed the experiment alone. In group decision-making, however, confidence plays an important role. Research indicates that decision-makers automatically communicate confidence (45) with opinions expressed with higher confidence gaining more weight (46), and multiple studies detail how group members unify different expressions of individual confidence and cope with individual biases in confidence (47, 48). Thus, it would be interesting to see how a manipulation of prior beliefs (inducing under- or overconfidence) would change both the content and extent of confidence sharing among participants in scenarios of joint decision-making. Particularly reports of having made the wrong response, likely reflecting post-decision evidence accumulation (18, 49), could be very informative and useful in a social context.

## Conclusion

We demonstrated that a manipulation of prior beliefs in task performance, either via comparative feedback or via changes in task difficulty, causally influences subsequent ratings of decision confidence for perceptual decisions. These findings were well accounted for within a dynamic probabilistic model of decision confidence by changing the model’s prior belief. Our findings provide a mechanistic understanding of under- and overconfidence.

## Materials and Methods

### Participants

Fifty participants (eight men, one third gender, age: M = 19, SD = 4.9, range 17–52) took part in Experiment 1. Fifty participants (five men, age: M = 18.5, SD = 1, range 17–22) took part in Experiment 2. Due to chance level performance in at least one of the tasks, we removed two participants from Experiment 1 and three from Experiment 2. All participants participated in return for course credit and read and signed a written informed consent at the start of the experiment. All procedures were approved by the local ethics committee.

### Stimuli and apparatus

Both experiments were conducted on a 22-inch DELL monitor with a 60 Hz refresh rate, using PsychoPy3 (50). All stimuli were presented on a black background centered around the middle of the screen (radius 2.49° visual arc). Stimuli for the dot number task (white dots) were presented in two equally sized boxes (height 20°, width 18°) at an equal distance from the center of the screen. Stimuli for the letter discrimination task (white X’s and O’s) and dot color task (red and blue dots) were presented in one box (height 22°, width 22°), centered around the fixation point.

### Procedure

#### General

In both experiments, participants completed three decision-making tasks: a dot color task, a dot number task and a letter discrimination task (see **Figure 2**). Each task started with 120 training trials. In Experiment 1, participants were presented performance feedback every 24 trials, while in Experiment 2, feedback was given on every trial. After the training phase of a task, a test phase of 216 trials followed during which no feedback was provided, but instead participants indicated their level of confidence after each choice. For all tasks, a trial started with a fixation cross that was presented for 500 ms, after which the stimulus appeared for 200 ms or until a response was given. Participants indicated their choice using the C or N key using the thumbs of both hands. There was no time limit for responding. On test trials, participants additionally rated their confidence after each choice on a 6-point scale, labeled ‘certainly wrong’, ‘probably wrong’, ‘maybe wrong’, ‘maybe correct’, ‘probably correct’, and ‘certainly correct’ (reversed order for half the participants). Confidence was indicated using the 1, 2, 3, 8, 9 and 0 keys with the ring, middle and index fingers of both hands. There was no response limit for indicating confidence.

For each task, there were three levels of stimulus difficulty (easy, average or difficult). Stimulus properties for Experiment 1 were decided based on the results of a small pilot study (N=5). For Experiment 2, stimulus properties were revised based on the results of Experiment 1 in order to achieve a better matching of accuracy between tasks. Stimulus dependencies for each task can be found in **Table 1**.

**Table 1.** Stimulus properties for each difficulty level, task and experiment.

|  |  | **Dot color task** |  | **Dot number task** |  | **Letter discrimination task** |  |
| --- | --- | --- | --- | --- | --- | --- | --- |
|  |  | *Number of dominant color dots (80 dots in total).* |  | *Number of dots in the variable field (reference field contains 50).* |  | *Number of dominant letters (80 letters in total)* |  |
|  |  | **EXP 1** | **EXP 2** | **EXP 1** | **EXP 2** | **EXP 1** | **EXP 2** |
| **Difficulty** | Easy | 61 – 65 | 61 – 65 | + or - 21 – 25 | + or - 21 – 25 | 70 – 75 | 70 – 74 |
|  | Average | 51 – 55 | 46 – 50 | + or - 11 – 15 | + or - 11 – 15 | 51 – 55 | 53 – 57 |
|  | Difficult | 41 – 45 | 41 – 45 | + or - 1 – 5 | + or - 1 – 5 | 41 – 45 | 42 – 46 |

#### Dot color task

On each trial, participants decided whether a field contained more (static) blue or red dots. The total number of dots was always 80, with differing proportions of red or blue dots depending on the difficulty condition. The position of dots was randomly generated on each trial.

#### Dot number task

On each trial, two fields were presented, one of which contained 50 dots and the other more or less than 50 dots. Participants decided which of the two fields contained the largest number of dots. The exact number of dots in the variable field differed depending on the difficulty condition. The position of dots was randomly generated on each trial.

#### Letter discrimination task

On each trial, participants decided whether a field contained more X’s or O’s. The total number of X’s and O’s was always 80, with differing proportions of X’s or O’s depending on the difficulty condition. The position of the letters was randomly generated on each trial.

### Experiment 1: Prior belief induction in the comparative feedback experiment

In Experiment 1, prior beliefs about the ability to correctly perform the task were manipulated by means of fake comparative feedback during the training phase. Participants were told that their feedback score was indicative of their performance (accuracy and reaction time) on the preceding trials relative to the performance of other participants who took part previously. Unknown to participants, feedback was predetermined to be either good, average or bad for a specific task, and feedback scores were randomly sampled according to the feedback condition. Each participant received good feedback on one task (inducing prior beliefs of low task performance), average feedback on another task, and bad feedback on a third task (inducing prior beliefs of high task performance; order and mapping with tasks counterbalanced between participants). For each task, participants received feedback after every 24 training trials, amounting to 5 feedback presentations per task. Feedback scores were pseudo-randomly generated on each feedback presentation and ranged between 5 and 30% in the negative feedback condition, between 37 and 62% in the average feedback condition and between 70 and 95% in the positive feedback condition. To increase credibility of the negative feedback, the second out of the five feedback screens showed average feedback (ranging between 32% and 36%, labeled as average). Likewise, the second out of five feedback screens in the positive feedback condition showed average feedback (ranging between 63% and 67%, labeled as average).

At the top of feedback screens, a verbal indication of the participant’s score was presented: depending on the score, “Good performance:” in green, “Average performance:” in white or “Bad performance:” in red. After the colon, the score itself was presented in the same color as the verbal indication. In the middle of the feedback screen, the participant’s score was indicated in a visual way. A vertically oriented rectangle with no fill color was presented, with the bottom line marked “worst performance”, the top line marked “best performance” and a midline marked “average performance”. The participant’s score was used to color the same percentage of the rectangle’s total surface (starting at the bottom) in red (bad performance), white (average performance) or green (good performance) (see **Figure 2**).

### Experiment 2: Prior belief induction via task difficulty

In Experiment 2, prior beliefs about the ability to correctly perform the task were induced by manipulating the difficulty of the task during the training phase in three levels. Contrary to Experiment 1, participants received genuine feedback on every trial: Each correct choice was followed by the word “Correct!” and each incorrect choice by “Wrong!”. Each participant completed one task with a training phase consisting of only easy trials (inducing positive prior beliefs about task ability), another with a training phase of all average trials (inducing average prior beliefs), and another with a training phase of all difficult trials (inducing negative prior beliefs).

### Statistical analyses

Data from the test phase were analyzed using mixed effects models. We started from models including the fixed factors testing phase difficulty and condition (Experiment 1: positive, average or negative feedback; Experiment 2: easy, average or hard training phase) and their interaction, as well as a random intercept for each participant. These models were then extended by adding random slopes, only when this significantly improved model fit. For Experiment 1, the final model for confidence ratings included additive slopes of both testing phase difficulty and feedback, while the final models for accuracy and RTs included only a random slope of feedback. For Experiment 2, all three models included only a random slope of training condition. Confidence ratings and reaction times were analyzed with linear mixed effects models, for which we report *F* statistics and the degrees of freedom as estimated by Satterthwaite’s approximation. Accuracy was analyzed using a generalized linear mixed model, for which we report *χ*^2^ statistics. All model fit analyses were done using the lmerTest package (51) in RStudio (52). In addition to these frequentist analyses, we calculated Bayes factors (BF) using the BayesFactor package in R (53) with default priors. A BF_10_ indicates data in favor of the null hypothesis (BF_10_ < 1/3), data in favor of the alternative hypothesis (BF_10_ > 3), and data that is uninformative (BF_10_ ≈ 1).

### Computational model

#### Bounded evidence accumulation

We modeled the data using the drift diffusion model (DDM), a popular variant of the wider class of accumulation-to-bound models. In the DDM, noisy evidence is accumulated, the strength of which is controlled by a drift rate *v*, until one of two boundaries *a* or *-a* is reached. Non-decision components are captured by a non-decision time *ter* parameter. To simulate data from the model, random walks were used as a discrete approximation of the continuous diffusion process of the drift diffusion model (54). Each simulated random walk process started at *z*\**a* (here, *z* was an unbiased starting point of 0) which terminated once the accumulated evidence reached either *a* or -*a*. At each time step *τ*, accumulated evidence changed by Δ with Δ given in Eq. (1):

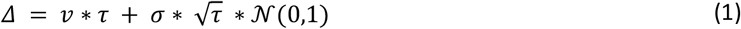

Within-trial variability is given by *σ*. In all simulations, *τ* was set to 1 ms, and *σ* was fixed to .1.

#### Accounting for prior beliefs

Within this model, confidence is given by mapping accumulated evidence, reaction time and the choice on a 2-dimensional heatmap (as shown in **Figure 1**) representing the probability of being correct for any given evidence level, time and choice. Since confidence judgments were given after the choices in both experiments, we allowed for additional post-decision evidence accumulation following boundary crossing before quantifying confidence (49). The duration of the post-decision evidence accumulation process was sampled from the full confidence reaction time distribution observed during the test phase for each subject. The heatmaps were constructed by computing the ratio between the probability densities of the amount of evidence accumulated with a given drift rate *μ* > 0 and its opposite -μ at each time step (the inverse ratio is computed depending on the choice). An important aspect is that these heatmaps depend on the actual drift rate that is used to generate them; when generating heatmaps with high versus low drift rates, the probability of being correct will be high versus low, respectively (because high drift rates are associated with a higher accuracy and vice versa). To model prior beliefs, we assumed that the drift rate parameter controlling the shape of the heatmap can be different from the drift rate parameter controlling objective performance. To avoid confusion, we refer to the former as the subjective drift rate (*v*_*s*_, formalizing the theoretical notion of prior beliefs) and the latter as the drift rate (*v*).

#### Qualitative model fitting

We estimated *v*_*s*_ for each participant and each prior belief condition by estimating which *v*_*s*_ provides predictions about confidence that best match the feedback received by participants in the training phase. Note that in Experiment 1, feedback was only given once at the end of each training block (24 trials), so we equally assigned the feedback value presented at the end of a block to every trial within that block. To have access to the amount of accumulated evidence, we first simulated predictions of the observed trials in the training phase from DDM parameters fitted to the training data. Confidence predictions for those simulated trials were then quantified as the probability correct given time and evidence for the heatmap generated by *v*_*s*_. The cost function was determined by the mean squared error (MSE) between observed feedback and predicted confidence Eq (2):

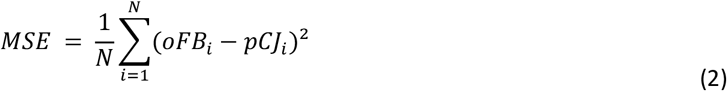

with *N* the number of observed trials in the training phase for a given prior belief condition, *oFB*_*i*_ the feedback received at trial *i* and *pCJ*_*i*_ the confidence predicted for trial *i*. Each observed trial’s feedback was compared 24 times to new predictions to account for the stochastic nature of the DDM. Since generating a heatmap is computationally costly, we generated 500 heatmaps from values of v_s_ ranging from 0 to .5. The MSE was then computed for each of these generated heatmaps. Smoothing using the locally weighted scatterplot smoothing method (LOWESS; (55)) was performed over the computed MSE for all *v*_*s*_ to further reduce noise. The final estimated *v*_*s*_ for each participant and prior belief condition was therefore equal to the one that generated the heatmap with the minimum smoothed MSE.

#### Quantitative model fitting

Quantitative model predictions were produced by directly fitting our model to confidence ratings in the testing phase. An improved implementation of the heatmap generation allowed us to directly estimate the best-fitting *v*_*s*_, instead of comparing the cost for several pre-generated values like explained in previous section. We estimated v_s_ separately for each participant and prior belief condition. Since model confidence is given as a probability of being correct, we applied equal-width binning to map model predictions on the confidence ratings scale. The biases and individual differences in mapping confidence on a categorical scale were accounted for by estimating an additional bias parameter separately for each participant, but fixed over conditions. To estimate these parameters, we computed the proportion of trials falling in each confidence level, separately for corrects and errors. We then used a differential evolution algorithm, as implemented in the DEoptim R package (56), to minimize the sum of squared error function shown in Eq (3):

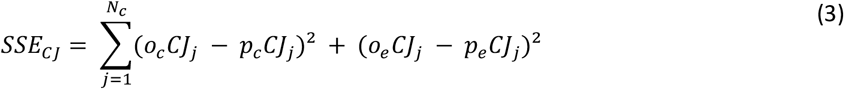

With *N*_*c*_ the number of confidence levels, *o*_*c*_*CJ*_*j*_ and *p*_*c*_*CJ*_*j*_ the proportion of observed and predicted correct trials with a confidence judgment *j*, respectively, and *o*_*e*_*CJ*_*j*_ and *p*_*e*_*CJ*_*j*_ the proportion of observed and predicted incorrect trials with a confidence judgment j. The population size for the differential evolution algorithm was set to 10 times the number of free parameters, as recommended in (57). Two termination criteria were set: (1) no new minimum of the SSE observed for the past 100 iterations or (2) a maximum of 1000 iterations. The 1000 iterations criterion was never reached. Model predictions for the sake of parameter estimation were generated by simulating 5000 random walk paths for each drift rate to be fitted. Model predictions from best-fitting parameters (as shown on **Figure 3**) were generated by simulating an equal amount of paths as the corresponding observed data.

#### DDM fitting

For each task and participant in the training data of Experiment 1 as well as in the test data of both experiments, we fitted 5 DDM parameters to the accuracy and reaction time data: 3 drift rates (*v*; one for each trial difficulty level), the decision boundary (*a*) and the non-decision time (*Ter*). Since only one trial difficulty was presented per task in the training phase of Experiment 2, only one drift rate per task was fitted to the training data of Experiment 2, resulting in the estimation of 3 DDM parameters in this case. To estimate these parameters, we implemented quantile optimization. Specifically, we computed the proportion of trials in six groups formed by quantiles .1, .3, .5, .7 and .9 of reaction time, separately for corrects and errors. We used a differential evolution algorithm to minimize the following SSE:

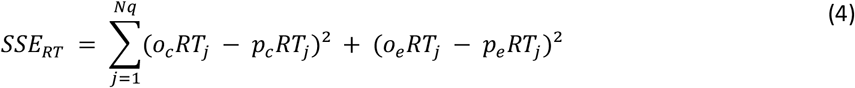

with *N*_*q*_ the number of quantiles, *o*_*c*_*RT*_*j*_ and *p*_*c*_*RT*_*j*_ the proportion of observed and predicted correct responses in RT quantile *j*, respectively, and *o*_*e*_*RT*_*j*_ and *p*_*e*_*RT*_*j*_ the proportion of observed and predicted incorrect responses in RT quantile *j*. Model fitting was done separately for each participant, phase (training vs testing), and experimental manipulation. All DEoptim settings were identical to the ones described in previous section.

## Acknowledgments

This research was supported by a Francqui start-up grant (PXF-D8830), a starting grant from the KU Leuven (STG/20/006) and a grant by the Research Foundation Flanders, Belgium (FWO-Vlaanderen) (G0B0521N), awarded to K.D., and a grant by the Research Foundation Flanders, Belgium (FWO-Vlaanderen) awarded to H.V.M. (11E6423N). All raw data and analysis code are openly available at [insert link upon publication].

## Supplementary materials

**Figure S1.**
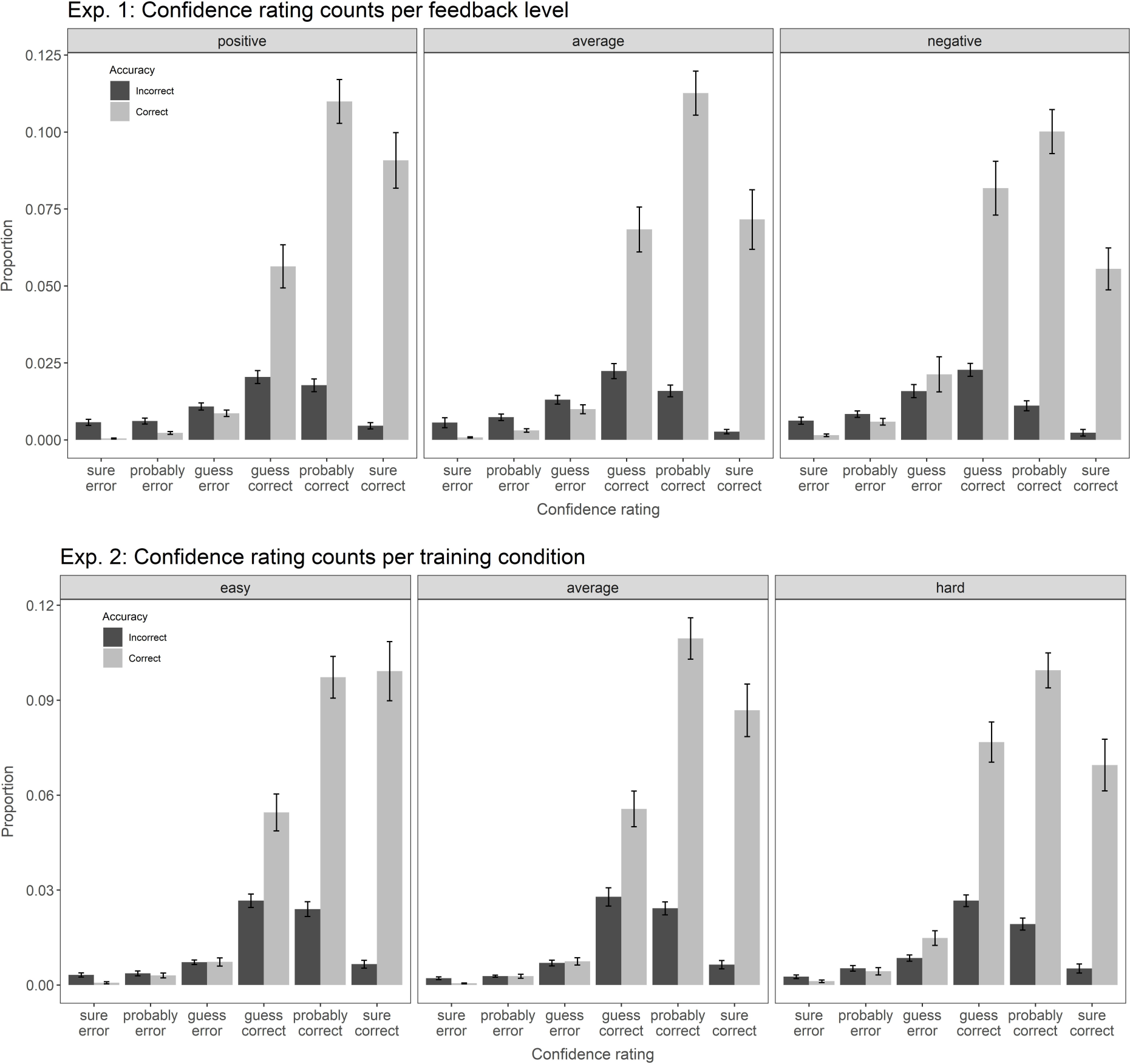
Confidence ratings in Experiment 2. Mean proportions of confidence ratings for correct and incorrect choices in Experiment 1 (top panel) and Experiment 2 (bottom panel), split per feedback level and training difficulty level, respectively. Error bars indicate standard errors.

**Figure S2.**
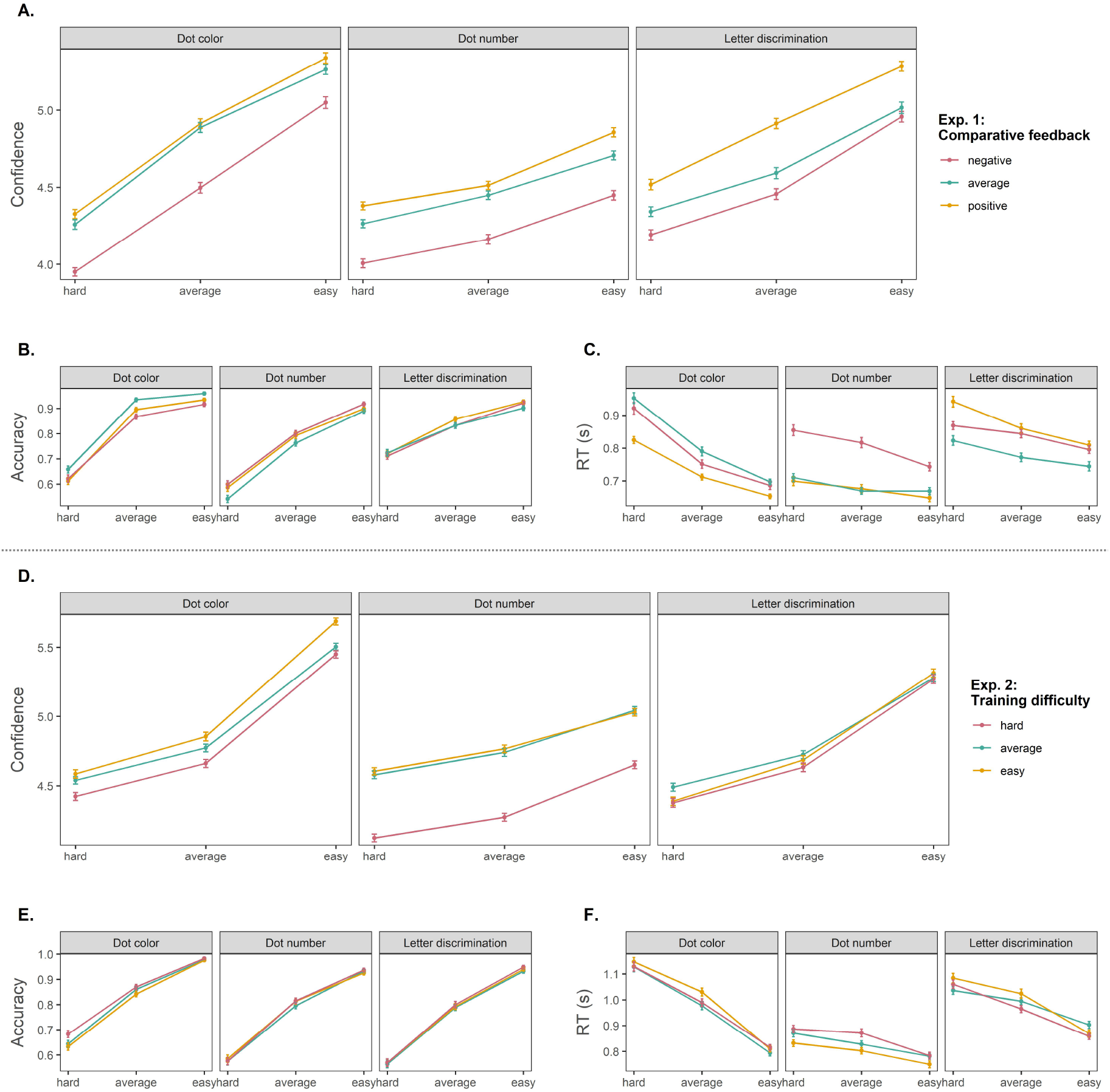
Effects of prior beliefs manipulations per task. In both experiments, three different tasks were used to assess the effect of three prior beliefs conditions: a dot color task, a dot number task and a letter discrimination task. The mapping between tasks and conditions, as well as their order, was counterbalanced. **(A-B)** Effect of three different levels of comparative feedback (negative, average or positive) on confidence ratings, accuracy and RTs, respectively, for each task separately. **(D-F)** Effect of three different levels of training difficulty (hard, average, easy) on confidence ratings, accuracy and RTs, respectively, for each task separately. Note: error bars reflect standard error of the mean.

